# RNF213 variant and autophagic impairment: A pivotal link to endothelial dysfunction in Moyamoya disease

**DOI:** 10.1101/2023.10.11.561969

**Authors:** Hee Sun Shin, Geun Hwa Park, Eun Sil Choi, So Young Park, Da Sol Kim, Jaerak Chang, Ji Man Hong

## Abstract

**Background:** Moyamoya disease (MMD) is closely associated with the Ring Finger Protein 213 (RNF213), a susceptibility gene for this disease. However, its biological function remains unclear. We aimed to elucidate the role of RNF213 in the damage incurred by human endothelial cells under oxygen-glucose deprivation (OGD), a condition that mimics intracranial ischemia in patients with MMD.

**Methods:** We analyzed autophagy in peripheral blood mononuclear cells (PBMCs) derived from patients carrying either RNF213 wild-type (WT) or variant (R4810K). Subsequently, human umbilical vein endothelial cells (HUVECs) were transfected with RNF213 WT (HUVEC^WT^) or R4810K (HUVEC^R4810K^) and exposed to OGD for 2 h to determine the role of the RNF213 variant in such a setting. Immunoblotting was used to analyze autophagy marker proteins, and tube formation assays were performed to examine endothelial function. Autophagic vesicles were observed using transmission electron microscopy. Post-OGD exposure, we administered autophagy modulators such as rapamycin and cilostazol.

**Results:** The RNF213 variant group during post-OGD exposure (vs. pre-OGD exposure) showed autophagy inhibition, increased protein expression of SQSTM1/p62 (*p* < 0.0001) and LC3-II (*p* = 0.0039), and impaired endothelial function (*p* = 0.0252). HUVEC^R4810K^ during post-OGD exposure (versus pre-OGD exposure) showed a remarkable increase in autophagic vesicles. Administration of autophagy modulators notably restored the function of HUVEC^R4810K^ and cellular autophagy.

**Conclusions:** Our findings support the pivotal role of autophagy impaired by the RNF213 variant in MMD-induced endothelial cell dysfunction and underscore the critical mechanism of autophagy leading to progressive endothelial dysfunction and MMD pathogenesis under relative ischemia within the intracranial portion.

## Introduction

Moyamoya disease (MMD) is an intracranial vascular disorder characterized by progressive specialized occlusive lesions of the terminal internal carotid arteries (ICA). ^1, 2^ Endothelial dysfunction is significantly increased in MMD patients. ^3, 4^ Despite the fact that most MMD patients are sporadic, familial MMD is present in approximately 15% of East Asian countries such as Japan, Korea, and China. ^5, 6^ Revascularization surgery is the main treatment strategy for MMD patients^7^, and antiplatelets can be another option to prevent clinical deterioration in MMD patients. ^8, 9^

In 2011, as part of a genome-wide association study (GWAS), Ring Finger Protein 213 (RNF213) was identified as a susceptibility gene for MMD. ^2, 10^ Although the function of RNF213 is not entirely clear, previous studies have suggested that it mainly plays a role in angiogenesis and the function of vascular endothelial cells. ^11, 12^ The most commonly found variant of RNF213 in MMD patients is R4810K (rs112735431, c.14429G>A), which is frequently identified in East Asia. ^10, 13, 14^ This variant has been associated with earlier onset and more severe deterioration of MMD. ^15–18^ The heterozygous RNF213 R4810K variant is more prevalent in adult patients with MMD and is associated with a higher risk of cerebral hemorrhage, whereas the homozygotic variant is commonly found in children with MMD. ^19^

Structurally, RNF213 is a large protein, measuring 591kDa, which contains AAA+ ATPase and E3 ligase domains. ^2^ Variants associated with MMD were predominantly observed in the E3 ligase domain. ^20^ E3 ligase plays a central role in ubiquitination by recruiting an E2 ubiquitin-conjugating enzyme to attach ubiquitin to its substrate. Ubiquitin serves as a signaling molecule that controls a wide range of biological processes, including protein degradation, autophagy, immunity, and inflammation. RNF213 has also Ring finger (RING) domain that cooperates with the Ubc13 E2 ubiquitin-conjugating enzyme. The RNF213 R4810K variant is ubiquitinated. ^21^ Thus, it is believed that the reduced ubiquitination caused by this variant leads to the accumulation of waste substrates.

Autophagy is a dynamic process that maintains normal physiological conditions. ^22^ It promotes cell survival by degrading long-lived and misfolded proteins and damaged organelles to generate amino acids for energy metabolism. This process is particularly important under stressful conditions such as starvation or hypoxia. ^23^

Some studies have investigated diabetic and atherogenic conditions, which are high-risk factors for MMD. ^24, 25^ In addition, advanced glycation end-products (AGEs), which contribute to the pathogenesis of diabetes mellitus (DM) and atherosclerosis, induce autophagy through the production of reactive oxygen species (ROS) in human umbilical vein endothelial cells (HUVECs). ^26^ In patients with DM, autophagy is impaired in endothelial cells. ^27^ Additionally, autophagy was impaired by tumor necrosis factor (TNF)-induced inflammation in HUVECs. ^28^ Previous studies have shown that inhibition of autophagy enhances endothelial cell injury in response to various external factors (e.g., OGD, inflammation, and DM). ^27–29^ Therefore, restoration of autophagy may be an effective treatment for a range of vascular diseases, including MMD. In this study, we investigated the possibility that the RNF213 R4810K variant is associated with endothelial dysfunction due to impaired autophagy caused by ischemia.

## Materials and methods

### Patients

Patients with MMD aged between 19 and 80 years were prospectively recruited from the Department of Neurology at Ajou University Hospital, Suwon, Republic of Korea. The selection criteria were: (1) subjects carrying wild-type RNF213 or (2) subjects carrying the RNF213 R4810K variant. Among the non-MMD group, patients with headache, dissection, vasculitis, and evidence of cardio-embolism were included as controls. This study adhered to the Declaration of Helsinki and was approved by the Institutional Review Board of Ajou University Medical Center (BMR-SMP-21- 357). Written informed consent was obtained from all participants.

### Flow mediated dilation (FMD)

Endothelial cellular function was evaluated clinically by measuring the brachial artery’s FMD in response to reactive hyperemia. The participants rested fully before the FMD measurements. A blood pressure cuff was placed on the subject’s forearm and inflated to 50 mmHg above the systolic pressure for 5 min to create an occlusion. The cuff pressure was promptly released to induce reactive hyperemia in the hand and forearm, leading to subsequent reactive vasodilation of the brachial artery. FMD was calculated as the maximum percentage change in the diameter of the brachial artery after reactive hyperemia compared with baseline.

### Isolation of human peripheral blood mononuclear cells (PBMCs)

Peripheral blood (8 ml) was collected from patients carrying either the RNF213 WT or R4810K into a CPT mononuclear cell preparation tube (362761, BD Biosciences, Franklin Lakes, NJ, USA). PBMCs were isolated according to the manufacturer’s instructions. Peripheral blood was collected in CPT tubes, which were then centrifuged for 10 min at 3,200 rpm. The plasma section was removed, and the PBMCs layer was collected. The PBMCs were cultured in RPMI-1640 medium (10-040-CV, Corning, Bedford, MA, USA) supplemented with 10% fetal bovine serum (FBS) and 1% penicillin-streptomycin and incubated in a humidified incubator at 37 ℃ and 5% CO2. To induce in vitro ischemia, PBMCs were washed with 1X DPBS (Lonza, Basel, Switzerland), switched to serum-and glucose-free medium (Gibco, Thermo Fisher Scientific Ltd., Waltham, MA, USA), and placed in a hypoxic chamber (Thermo Scientific Ltd., Waltham, MA, USA) for 2 h. The chamber was sealed and maintained in a gas mixture containing 85% N2 and 5% CO2. After 2 h of oxygen-glucose deprivation (OGD), PBMCs were returned to standard culture conditions in a complete RPMI-1640 medium containing 10% FBS and 1% penicillin-streptomycin for 1 h.

### Human umbilical vein endothelial cells (HUVECs) culture and treatments

HUVECs (Lonza, Basel, Switzerland) were cultured in an endothelial growth medium bullet kit (EGM- 2; Lonza, Basel, Switzerland) and used for experiments between passages three and seven. Cells were maintained at 37 °C in a 5% CO2 humidified atmosphere, with the culture medium replaced every other day. When the cells reached 70% confluence after transfection, they were treated with rapamycin (RA, Sigma-Aldrich, Saint Louis, MO, USA) (40 nM) for 20 h or cilostazol (CSZ, Sigma-Aldrich, Saint Louis, MO, USA) (30 μM) for 24 h. The vehicle group was treated with 1% dimethyl sulfoxide.

### OGD condition

HUVECs were cultured in a complete medium (EGM-2) under standard culture conditions. To induce OGD, HUVECs were washed with 1X DPBS (Lonza, Basel, Switzerland), switched to serum-and glucose-free no-glucose medium (Gibco, Thermo Fisher Scientific Ltd., Waltham, MA, USA), and placed in a hypoxic chamber (Thermo Scientific Ltd., Waltham, MA, USA) for 2 h. The chamber was sealed and maintained in a gas mixture containing 85% N2 and 5% CO2. OGD stimulation was terminated by replacing the medium with a complete medium (EGM-2) under standard culture conditions for 1 h.

### Plasmids and transfection

The RNF213 full clone in the pcDNA3.1 vector was purchased from Genescript (Clone ID: OHu28328D). Bioneer generated a single variant of RNF213, R4810K (c.14576G>A). For transient overexpression, HUVECs were transfected with 2 μg of RNF213 WT or R4810K construct using electroporation (Lonza, Basel, Switzerland), following the manufacturer’s protocol.

### Western blot

A general procedure was used for western blot analysis. HUVECs were rinsed with 1X DPBS (Lonza, Basel, Switzerland) and lysed in RIPA buffer (MB-077-0050, Rockland immunochemicals, Philadelphia, PA, USA). The samples were centrifuged for 10 min at 13,000 rpm, 4 °C, and the supernatants were used for western blot. Protein concentrations were measured using a Protein Assay Kit (Thermo Scientific Ltd., Waltham, MA, USA) with bovine serum albumin as the standard. The samples (30 μg protein/well) were subjected to 5% or 13% sodium dodecyl sulfate-polyacrylamide gel electrophoresis (SDS-PAGE) and transferred to polyvinylidene difluoride membranes (PVDF, Millipore, Bedford, MA, USA). The following primary antibodies were used in this study: RNF213 (sc-293391, Santa Cruz Biotechnology, Santa Cruz, CA, USA), SQSTM1/p62 (610832, BD Biosciences, Franklin Lakes, NJ, USA), LC3-a/b (L8918, Sigma-Aldrich, Saint Louis, MO, USA), GAPDH (2118S, Cell Signaling Technology, Beverly, MA, USA); secondary antibodies: rabbit IgG horseradish peroxidase-conjugated antibody (GTX213110-01, Genetex, Irvine, CA, USA) or mouse IgG horseradish peroxidase-conjugated antibody (PI-2000, Vector Laboratories, Burlingame, CA, USA). Immunoreactive bands were visualized using an enhanced chemiluminescence detection system (WF329297; Thermo Scientific Ltd., Waltham, MA, USA). Differences in protein expression were analyzed by measuring the band intensities using ImageJ software (National Institutes of Health, Bethesda, MD, USA).

### Tube formation assay

To prepare the Matrigel matrix, 250 μl of Matrigel (354234, Corning, Bedford, MA, USA) was added per well of a 24-well plate and solidified for 30 min at 37 ℃. HUVECs were seeded at a density of 5 × 10^4^ cells per well on the Matrigel-coated plate and incubated in a humidified cell incubator with 5% CO2 at 37 °C for 24 h. After incubation, tube formation was observed under a phase-contrast inverted microscope. The number of branch points and total tube lengths were measured using the ImageJ software (National Institutes of Health, Bethesda, MD, USA).

### Transmission electron microscopy (TEM)

HUVECs transfected with RNF213 wild-type (WT) or R4810K under OGD conditions were fixed in 2% glutaraldehyde solution (15720, Sigma-Aldrich, Saint Louis, MO, USA). After fixation, the samples were washed with 0.1 M cacodylate buffer (pH 7.4) and post-fixed with 0.1 M cacodylate buffer with osmium tetroxide at room temperature for 2 h. The samples were then dehydrated with 50–100% graded ethanol solutions, infiltrated with propylene oxide, and embedded in Epon (Polysciences, Inc, Warrington, PA, USA). All sections were double stained with 6% uranyl acetate (Electron Microscopy Sciences, Inc, Hatfield, PA, USA). Finally, all ultrathin sections were observed using TEM (Sigma 500, Carl Zeiss, Germany) at the three-dimensional immune system imaging core facility of Ajou University.

### Statistical analysis

All statistical analyses were performed using Prism version 10 (GraphPad Software Inc., San Diego, CA, USA). Data are expressed as mean ± standard deviation (SD). To analyze the data statistically, we performed a one-way analysis of variance (ANOVA) for multiple comparisons in each group. They were followed by Tukey’s post hoc test. Statistical significance was set at p < 0.05.

## Results

### An MMD patient with RNF213 R4810K variant who was transiently malnourished

A 32-year-old man initially presented to a neurology outpatient clinic in 2019 for the management of recurrent episodes of typical migraine headaches. Upon initial assessment, computed tomography angiography indicated a putative stenosis in the right middle cerebral artery. Transfemoral cerebral angiography was performed to substantiate these initial observations (**Visit 1**), a transfemoral cerebral angiography (TFCA). Genetic testing revealed the presence of an RNF213 single-nucleotide polymorphism, specifically the R4810K variant, involving the substitution of arginine with lysine at position 4810. During this initial phase, the arterial stenosis was not deemed severe enough to warrant immediate intervention.

Consequently, the patient was managed on an outpatient basis and received pharmacological treatment for episodic migraines as needed. Approximately one year after the initial evaluation (in 2020), the patient experienced an abrupt deterioration in mental status, requiring immediate hospitalization. The patient was admitted with his father, who reported that he had run away from home several weeks prior and suffered from mental illness and malnutrition until law enforcement intervened. Clinical assessment revealed that the patient was lucid but had impaired peripheral vision and could not provide coherent responses during the threat test. He was wheelchair-dependent for mobility and had residual homonymous hemianopia. Based on these findings, subsequent hospitalization was initiated. Follow-up CT angiography (**Visit 2**) indicated a marked reduction in cerebral blood flow compared with the initial CT angiography. Additionally, TFCA revealed characteristic features of MMD, including pronounced shrinkage and stenosis in intracranial cerebral vessels, as well as bilaterally diminished cerebral blood flow, compared to prior findings (Figure 1).

**Figure 1.**
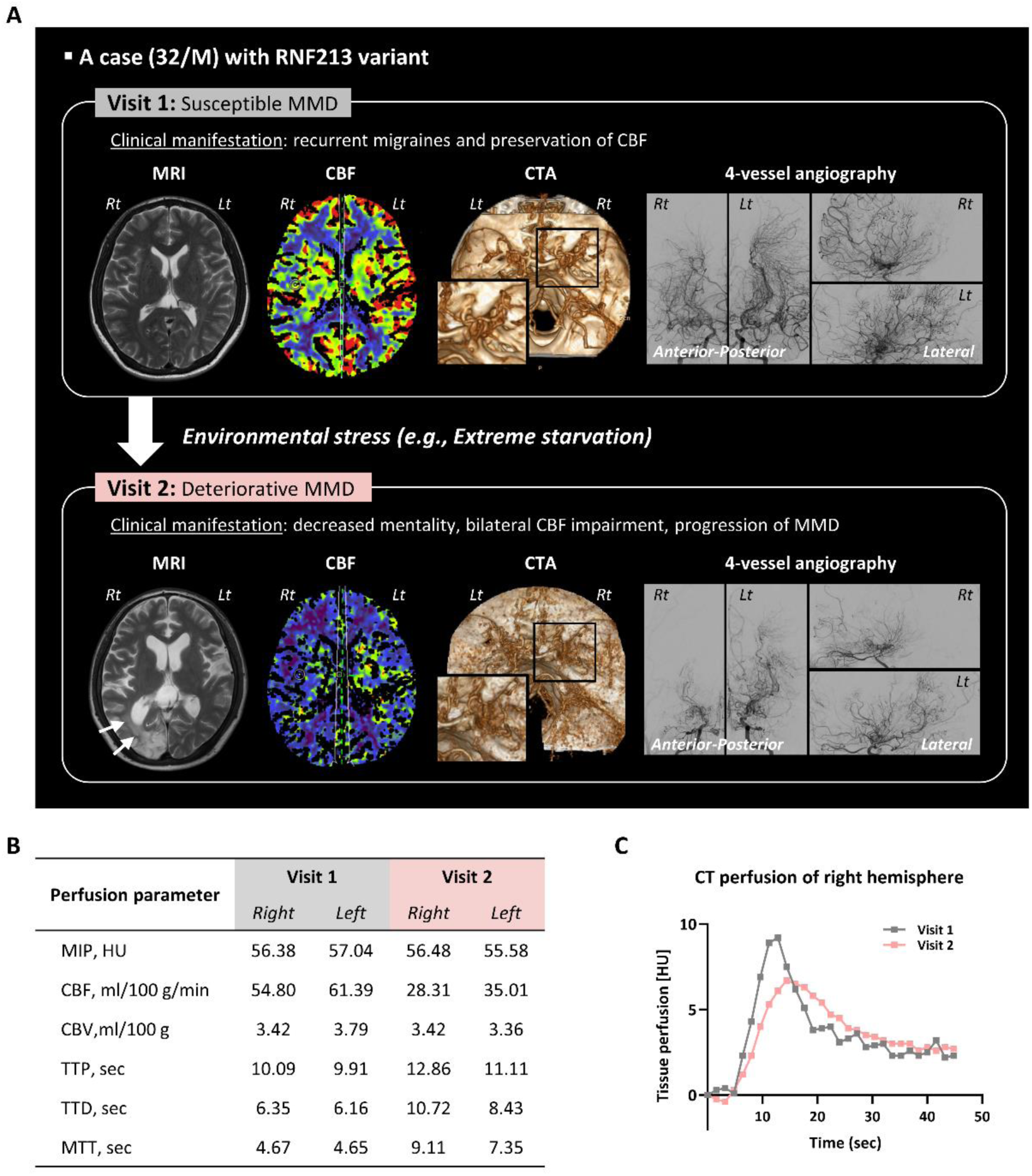
Sudden worsening of an MMD patient with RNF213 R4810K variant. (A) Depiction of clinical deterioration in a 32-year-old MMD patient with the RNF213 R4810K variant. Representative images (MRI, CBF, CTA, and angiographic findings) of the patient. The top panel showcases images from Visit 1, while the bottom panel presents images from Visit 2. The MRI from Visit 2 reveals cerebral infarction in the right posterior hemisphere (highlighted with a white arrow). CTA illustrates remarkable shrinkage of cerebral vessels (emphasized within a black square). CBF from Visit 2 shows a decrease compared to Visit 1. Transfemoral cerebral angiography (TFCA) images provide anterior, posterior, and lateral views of the anterior circulation. TFCA on Visit 2 also shows a considerable reduction of cerebral vessels on Visit 1. (B) The mean values of CTP between Visit 1 (gray column) and Visit 2 (pink column). (C) The time attenuation curve of CTP at Visit 1 (gray line) and Visit 2 (pink line). MRI, magnetic resonance imaging; CBF, cerebral blood flow; CTA, computed tomography angiography; CTP, computed tomography perfusion; MIP, maximum intensity projection; HU, Hounsfield Unit; CBV, cerebral blood volume; TTP, time to peak; TTD, time to drain; MTT, mean transit time.

### Various roles of the RNF213 variant in response to environmental stress

Since RNF213 was first discovered in 2011, numerous studies have attempted to elucidate its function and mechanism of action; however, they remain unclear. The most prevalent genetic variant among East Asian patients, RNF213 R4810K ^5, 6^, underscores the role of genetic factors in MMD pathophysiology. Both environmental and genetic factors have been reported to contribute to MMD progression. As shown in Table 1, various experimental models have examined the functional response of RNF213 to environmental stress. ^11, 30–32^ In this framework, the RNF213 variant appears to be activated under inflammatory, angiogenic, and hypoxic stress, leading to cellular dysfunction. In line with previous reports, we noted that clinical cases carrying the RNF213 R4810K variant evolved into MMD deterioration when exposed to intense environmental stressors, as exemplified by experimental oxygen-glucose deprivation. This prompted us to investigate the biological mechanisms of the RNF213 R4810K variant in relation to environmental triggers.

**Table 1.**
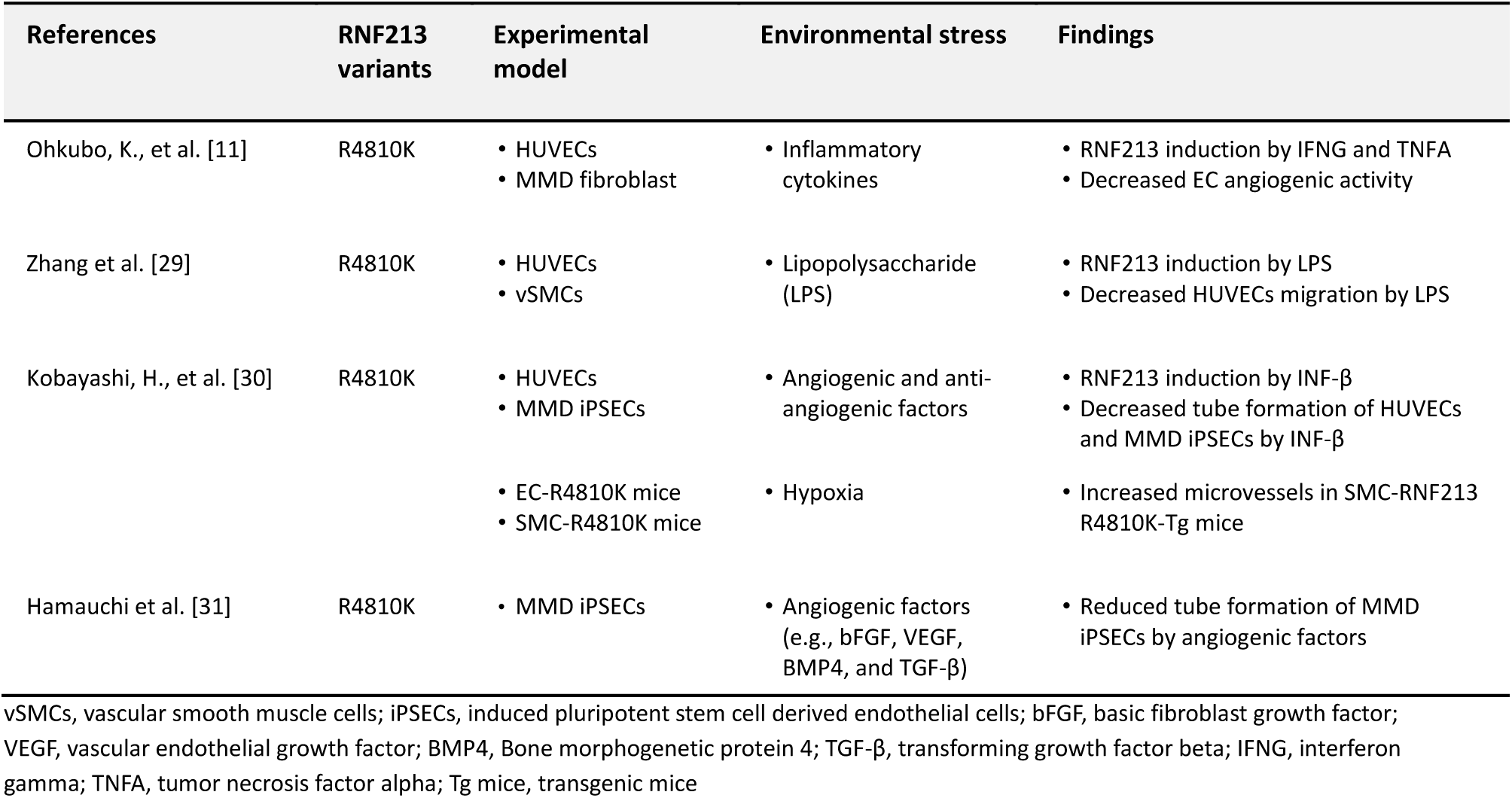
Overview of previous studies utilizing various experimental models related with RNF213 variant.

| References | RNF213 variants | Experimental model | Environmental stress | Findings |
| --- | --- | --- | --- | --- |
| Ohkubo, K., et al. [11] | R4810K | HUVECs<br>MMD fibroblast | Inflammatory cytokines | RNF213 induction by IFNG and TNFA<br>Decreased EC angiogenic activity |
| Zhang et al. [29] | R4810K | HUVECs<br>vSMCs | Lipopolysaccharide (LPS) | RNF213 induction by LPS<br>Decreased HUVECs migration by LPS |
| Kobayashi, H., et al. [30] | R4810K | HUVECs<br>MMD iPSECs | Angiogenic and anti-angiogenic factors | RNF213 induction by INF-β<br>Decreased tube formation of HUVECs and MMD iPSECs by INF-β |
|  |  | EC-R4810K mice<br>SMC-R4810K mice | Hypoxia | Increased microvessels in SMC-RNF213 R4810K-Tg mice |
| Hamauchi et al. [31] | R4810K | MMD iPSECs | Angiogenic factors (e.g., bFGF, VEGF, BMP4, and TGF-β) | Reduced tube formation of MMD iPSECs by angiogenic factors |

vSMCs, vascular smooth muscle cells; iPSECs, induced pluripotent stem cell derived endothelial cells; bFGF, basic fibroblast growth factor; VEGF, vascular endothelial growth factor; BMP4, Bone morphogenetic protein 4; TGF- $\beta$ , transforming growth factor beta; IFNG, interferon gamma; TNFA, tumor necrosis factor alpha; Tg mice, transgenic mice

### Autophagic impairment in MMD patients according to RNF213 R4810K variant

The genotype RNF213 R4810K was determined in consecutive admitted patients who presented with MMD. The patients were randomly assigned into 15 RNF213 wild-type (WT) and 15 RNF213 R4810K variant (R4810K) groups. To investigate whether there was a difference in endothelial cell function in MMD patients based on genotype (WT or R4810K), we measured FMD in MMD patients. FMD levels in the RNF213 R4810K group were significantly lower than those in the non-MMD group (Fig 2A).

**Figure 2.**
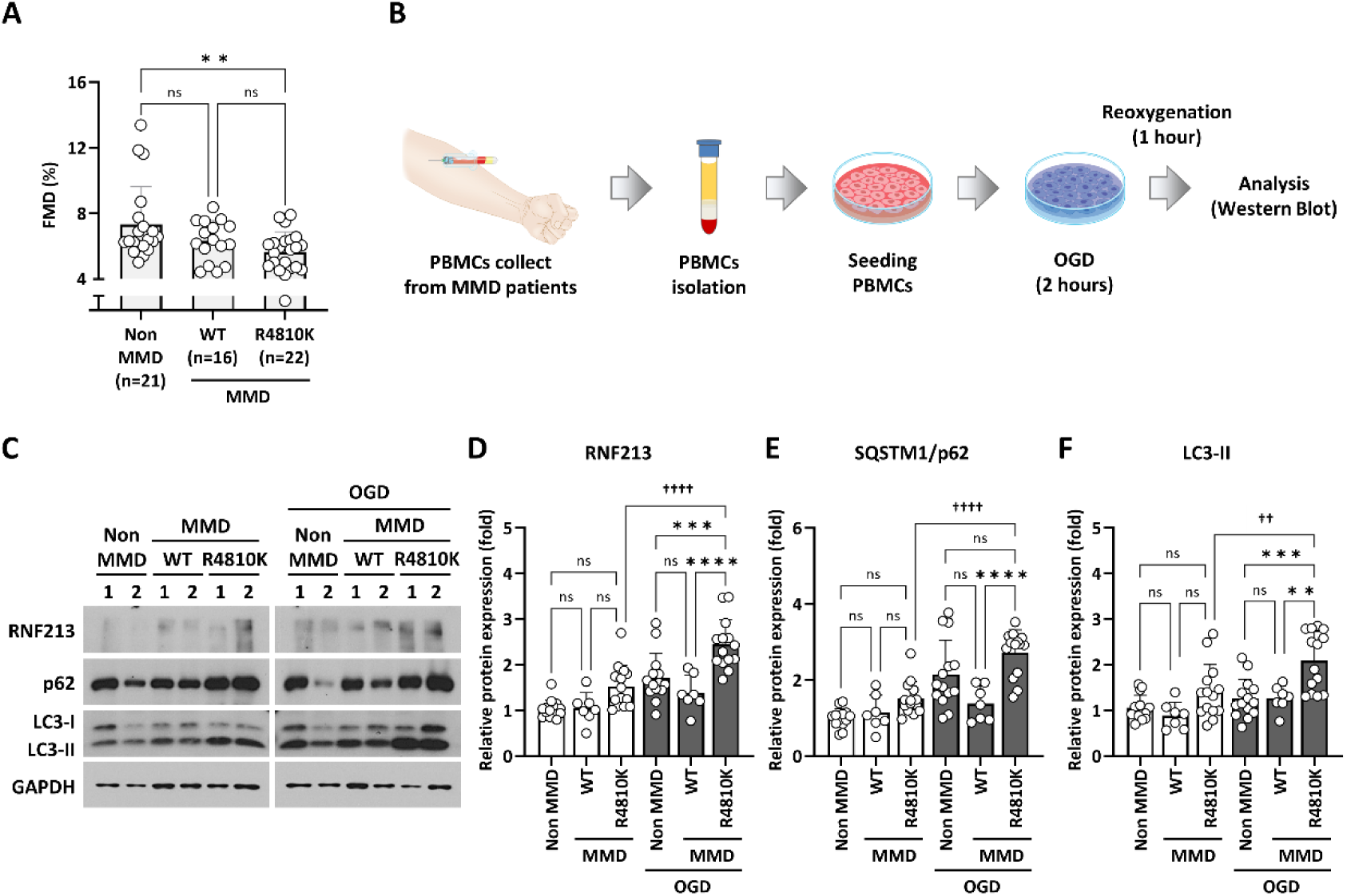
Impact of RNF213 R4810K variant on endothelial function and autophagy under oxygen-glucose deprivation (OGD) from MMD patient-derived PBMCs. (A) Comparative analysis showing decreased flow-mediated dilation (FMD, %) in MMD patients with RNF213 R4810K variant versus non-MMD patients. \*\**p* < 0.01. (B) An illustrative diagram presenting the sequential experimental process. MMD patient-derived PBMCs underwent 2-h OGD exposure followed by immunoblotting. (C) Immunoblotting results displaying protein expression levels in MMD patient-derived PBMCs with either RNF213 WT or R4810K and non-MMD-derived PBMCs in the absence or presence of OGD exposure. (D–F) Quantification graphs depicting relative protein levels associated with autophagy: RNF213 (R4810K group in pre-OGD vs. R4810K group in post-OGD, *p* < 0.0001), SQSTM1/p62 (R4810K group in pre-OGD vs. R4810K group in post-OGD, *p* < 0.0001), LC3-II (R4810K group in pre-OGD vs. R4810K group in post-OGD, *p* = 0.0039) normalized with GAPDH. The RNF213 R4810K variant in MMD patient-derived PBMCs inhibits autophagy during OGD exposure. RNF213 WT vs. R4810K^⁎^ (asterisks); pre-OGD vs. post-OGD^†^ (daggers). ^⁎⁎^*p* < 0.01, ^⁎⁎⁎^*p* < 0.001, ^⁎⁎⁎⁎^*p* < 0.0001, ^†^*p* < 0.05, ^††^*p* < 0.01, ^††††^*p* < 0.0001.

**Figure 3.**
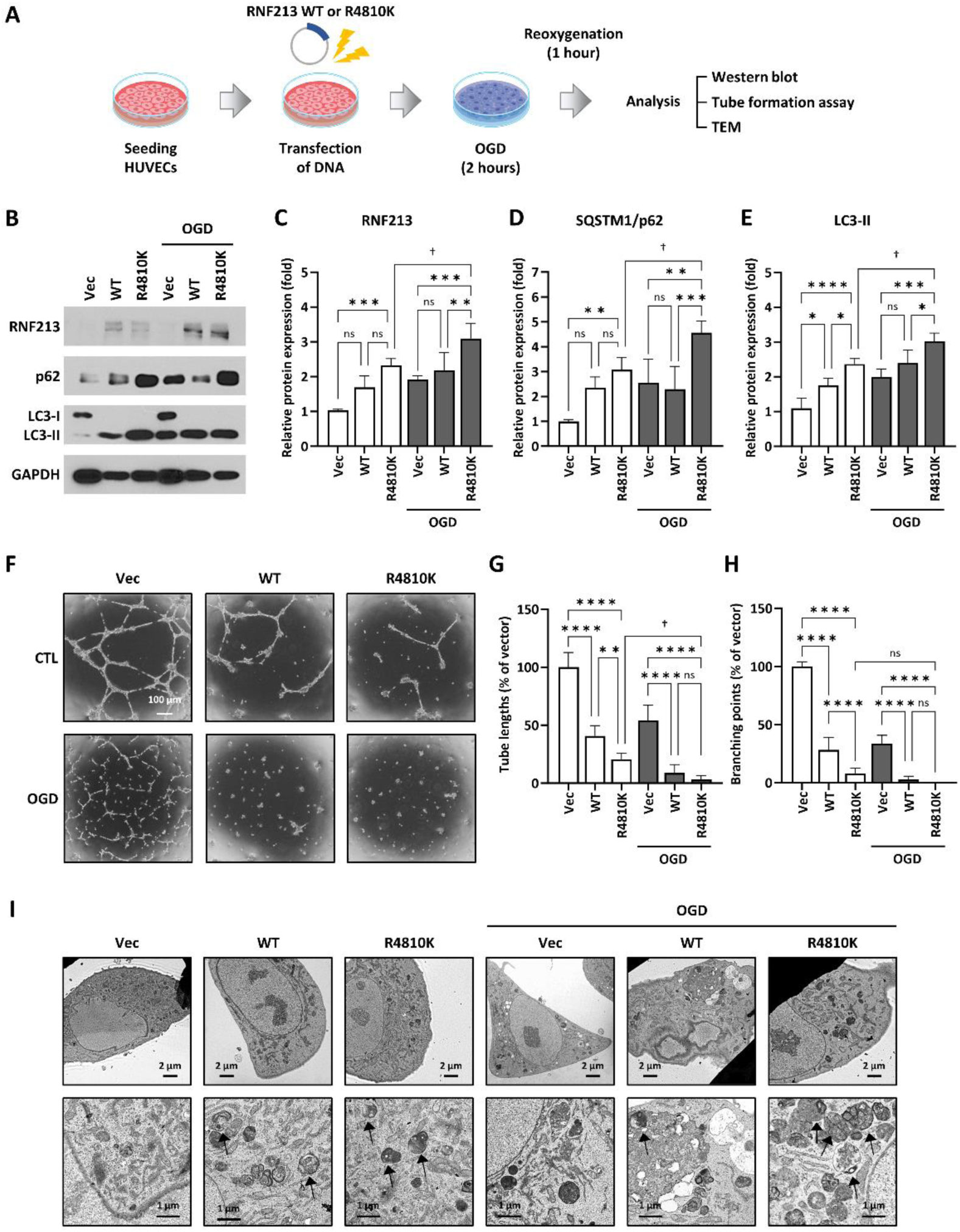
Impairment of autophagy in HUVECs transfected with RNF213 WT or R4810K under OGD exposure. (A) An illustrative diagram presenting the sequential experimental process. (B) Representative images of immunoblotting analysis for protein expression in transfected HUVECs with either empty vector (Vec), RNF213 WT, or R4810K under OGD exposure. (C–E) Quantification of relative protein expression related to autophagy: RNF213 (R4810K group in pre-OGD vs. R4810K group in post-OGD, *p* = 0.0274), SQSTM1/p62 (R4810K group in pre-OGD vs. R4810K group in post-OGD, *p* = 0.0366), LC3-II (R4810K group in pre-OGD vs. R4810K group in post-OGD, *p* = 0.0020) normalized with GAPDH (Values shown as mean ± standard deviation, based on three separate trials). (F) Representative images of tube formation assay are shown that reduced endothelial function in RNF213 R4810K variant group under OGD. Scale bar: 100 μm. (G and H) Quantification of tube lengths (G) and branching points (H) (Values shown as mean ± standard deviation, based on 3 separate trials). (I) Representative images of induced autophagic vesicles in RNF213 R4810K variant under OGD exposure, as assessed by transmission electron microscopy (TEM). Autophagic vesicles surrounded by a double membrane with undigested cytoplasmic material inside (black arrow). Scale bar: 2 μm (upper panels) and 1 μm (lower panels). RNF213 R4810K variant in HUVECs show autophagy inhibition and endothelial dysfunction during the experimental OGD exposure. RNF213 WT vs. R4810K^⁎^ (asterisks); pre-OGD vs. post-OGD^†^ (daggers). \*\**p* < 0.01, \*\*\**p* < 0.001, \*\*\*\**p* < 0.0001, ^†^*p* < 0.05.

To determine whether the autophagy function in PBMCs of MMD patients carrying the RNF213 WT or R4810K variant was affected by OGD exposure, we investigated the protein expression levels of RNF213, SQSTM1/p62, and LC3-II (Fig 2C). Notably, RNF213 expression was prominently increased in cells with the RNF213 R4810K variant following OGD exposure compared to before exposure (Fig 2D). The autophagy marker proteins SQSTM1/p62 and LC3-II expression were significantly higher in the PBMCs of patients post-OGD exposure as compared to pre-exposure (Fig 2E–F). These findings suggest that under cellular stress conditions, the RNF213 R4810K variant may suppress autophagy.

### Autophagic impairment in MMD patients according to RNF213 R4810K variant

The genotype RNF213 R4810K was determined in consecutive admitted patients who presented with MMD. The patients were randomly assigned into 15 RNF213 wild-type (WT) and 15 RNF213 R4810K variant (R4810K) groups. To investigate whether there was a difference in endothelial cell function in MMD patients based on genotype (WT or R4810K), we measured FMD in MMD patients. FMD levels in the RNF213 R4810K group were significantly lower than those in the non-MMD group (Fig 2A).

To determine whether the autophagy function in PBMCs of MMD patients carrying the RNF213 WT or R4810K variant was affected by OGD exposure, we investigated the protein expression levels of RNF213, SQSTM1/p62, and LC3-II (Fig 2C). Notably, RNF213 expression was prominently increased in cells with the RNF213 R4810K variant following OGD exposure compared to before exposure (Fig 2D). The autophagy marker proteins SQSTM1/p62 and LC3-II expression were significantly higher in the PBMCs of patients post-OGD exposure as compared to pre-exposure (Fig 2E–F). These findings suggest that under cellular stress conditions, the RNF213 R4810K variant may suppress autophagy.

### Rapamycin (RA)-induced restoration of autophagy and the function of HUVEC^R4810K^ under OGD exposure

We investigated the effects of RA on impaired autophagy in HUVECs transfected with RNF213 WT or R4810K variants. HUVEC^WT^ or HUVEC^R4810K^ were exposed to OGD for 2 h, followed by RA treatment for 20 h (Fig 4A). We found that when HUVECs were treated with RA, the upregulation of RNF213 protein levels by OGD exposure in HUVEC^R4810K^ was significantly decreased compared with that in the absence of OGD exposure (Fig 4B–C). Additionally, we confirmed that the expression of autophagy proteins (SQSTM1/p62 and LC3-II) induced by OGD exposure in HUVEC^R4810K^ was significantly decreased compared to that in the absence of OGD (Fig 4B, 4D, and 4E). We conducted a tube formation assay to assess functional changes in endothelial cells induced by RA. Under OGD conditions, RA significantly restored the impaired tube structure of HUVEC^R4810K^ cells (Fig 4F–H). Furthermore, we found that RA treatment recovered the autophagic vesicles induced in HUVEC^R4810K^ under OGD conditions (Fig 4I). Our data suggest that RA activates autophagy and recovers HUVECs expressing the RNF213 R4810K variant under OGD conditions.

**Figure 4.**
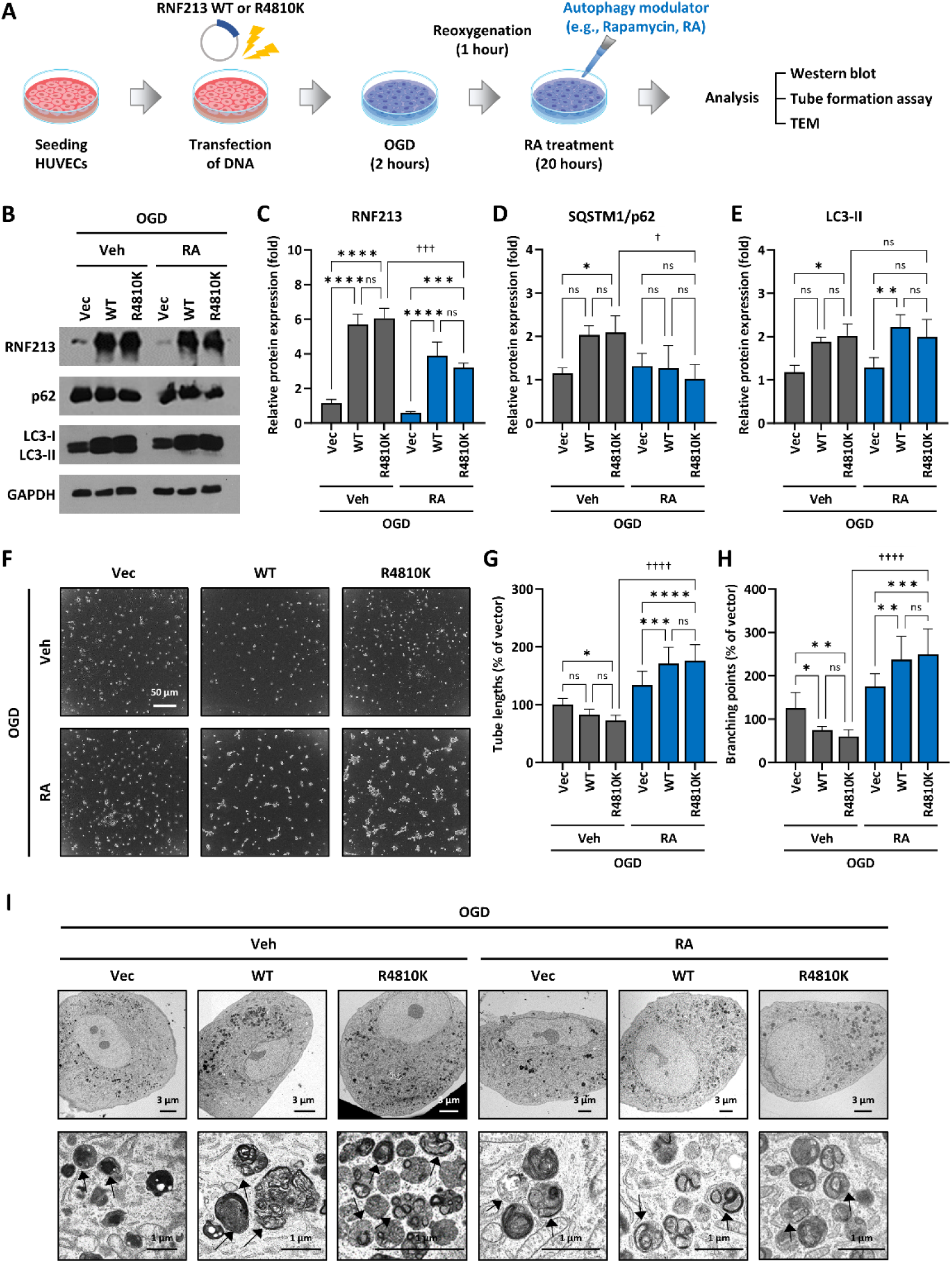
Restoration of suppressed autophagy by rapamycin (RA) in HUVECs transfected with RNF213 WT or R4810K under OGD. (A) An illustrative diagram of the experimental process for RA administration. (B) Representative images of immunoblotting analysis for protein expression in transfected HUVECs with either empty vector (Vec), RNF213 WT, or R4810K under OGD. (C–E) Quantification of relative protein expression related to autophagy: RNF213 (R4810K group treated with RA in pre-OGD vs. R4810K group treated with RA in post-OGD, *p* = 0.0001), SQSTM1/p62 (R4810K group treated with RA in pre-OGD vs. R4810K group treated with RA in post-OGD, *p* = 0.0187), LC3-II (R4810K group treated with RA in pre-OGD vs. R4810K group treated with RA in post-OGD, p>0.9999) normalized with GAPDH (Values shown as mean ± standard deviation, based on three separate trials). (F) Representative tube formation assay images show reduced endothelial function in the RNF213 R4810K variant group under OGD. Scale bar: 100 μm. (G and H) Quantification of tube lengths (G) and branching points (H) (Values shown as mean ± standard deviation, based on three separate trials). (I) Representative images of reduced autophagic vesicles by RA treatment in RNF213 R4810K variant under OGD, as assessed by transmission electron microscopy (TEM). Autophagic vesicles surrounded by a double membrane with undigested cytoplasmic material inside (black arrow). Scale bar: 3 μm (upper panels) and 1 μm (lower panels). Treatment of RA restores autophagy and endothelial function in HUVECs with RNF213 R4810K variant in response to OGD conditions. RNF213 WT vs. R4810K^⁎^ (asterisks); pre-OGD vs. post-OGD† (daggers) \**p* < 0.05, \*\**p* < 0.01, \*\*\**p* < 0.001, \*\*\*\**p* < 0.0001, ^†^*p* < 0.05, ^†††^*p* < 0.001, ^††††^*p* < 0.0001.

### Cilostazol (CSZ)-indued restoration of autophagy impairment and the function of HUVEC^R4810K^ under OGD exposure

We investigated the effects of CSZ on impaired endothelial cells because this drug is known to improve the symptoms of MMD. HUVEC^WT^ or HUVEC^R4810K^ were exposed to OGD for 2 h and then treated with CSZ for 24 h (Fig 5A). We found that when HUVECs were treated with CSZ, the upregulated RNF213 protein level under OGD exposure in HUVEC^R4810K^ cells was significantly decreased compared to that in the absence of OGD exposure (Fig 5B–C). Moreover, the induction of SQSTM1/p62 protein levels by OGD exposure in HUVEC^R4810K^ was remarkably reduced compared with that in the absence of OGD (Fig 5D). Treatment with CSZ significantly rescued the impaired tube formation of HUVEC^R4810K^ under OGD exposure (Fig 5F–H). Additionally, CSZ treatment recovered the autophagic vesicles upregulated by RNF213 WT or R4810K overexpression under OGD conditions (Fig 5I). Our data suggest that CSZ activates autophagy and restores HUVECs harboring the RNF213 R4810K variant under OGD conditions.

**Figure 5.**
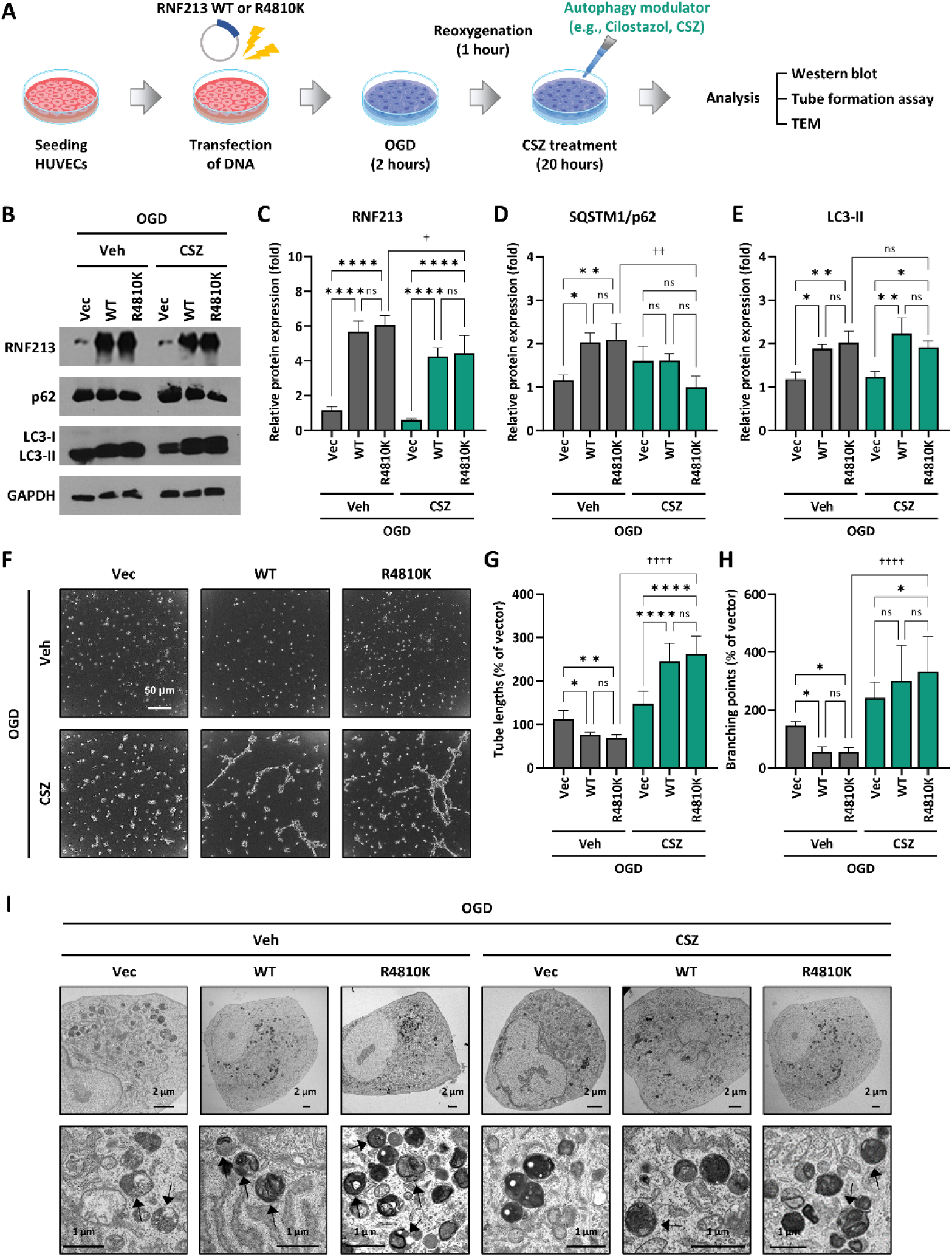
Restoration of suppressed autophagy by cilostazol (CSZ) in HUVECs transfected with RNF213 wild-type or R4810K under OGD. (A) An illustrative diagram of the experimental process for CSZ administration. (B) Representative images of immunoblotting analysis for protein expression in transfected HUVECs with either empty vector (Vec), RNF213 WT, or R4810K under OGD. (C–E) Quantification of relative protein expression related to autophagy: RNF213 (R4810K group treated with CSZ in pre-OGD vs. R4810K group treated with CSZ in post-OGD, *p* = 0.0484), SQSTM1/p62 (R4810K group treated with CSZ in pre-OGD vs. R4810K group treated with CSZ in post-OGD, *p* = 0.0026), LC3-II (R4810K group treated with CSZ in pre-OGD vs. R4810K group treated with CSZ in post-OGD, *p* = 0.9925) normalized with GAPDH (Values shown as mean ± standard deviation, based on three separate trials). (F) Representative tube formation assay images show reduced endothelial function in the RNF213 R4810K variant group under OGD. Scale bar: 50 μm. (G and H) Quantification of tube lengths (G) and branching points (H) (Values shown as mean ± standard deviation, based on three separate trials). (I) Representative images of reduced autophagic vesicles by CSZ treatment in the RNF213 variant under OGD, as assessed by transmission electron microscopy (TEM). Autophagic vesicles are surrounded by a double membrane with undigested cytoplasmic material inside (black arrow). Scale bar: 2 μm (upper panels) and 1 μm (lower panels). Treatment of CSZ restores autophagy and endothelial function in HUVECs with RNF213 R4810K variant in response to OGD condition. RNF213 WT vs. R4810K^⁎^ (asterisks); pre-OGD vs. post-OGD† (daggers) \**p* < 0.05, \*\**p* < 0.01, \*\*\*\**p* < 0.0001, ^†^*p* < 0.05, ^††^p < 0.01, ^††††^p < 0.0001.

## Discussion

Our research indicates that the RNF213 R4810K variant is sensitive to environmental stress and plays a significant role in the onset or development of MMD by adversely affecting autophagy. (Figure 6.) We observed a decline in endothelial function in patients with MMD carrying this specific variant. Notably, we observed significant arterial changes in the RNF213 R4810K variant, which causes malnutrition that triggers autophagy. Our in vitro analyses confirmed that the RNF213 R4810K variant substantially hampered the function of endothelial cells and was directly linked to the inhibition of autophagy. Impaired autophagy was restored by RA and CSZ (autophagy modulators), suggesting that this may be a potential treatment for patients with MMD harboring the RNF213 R4810K variant.

**Figure 6.**
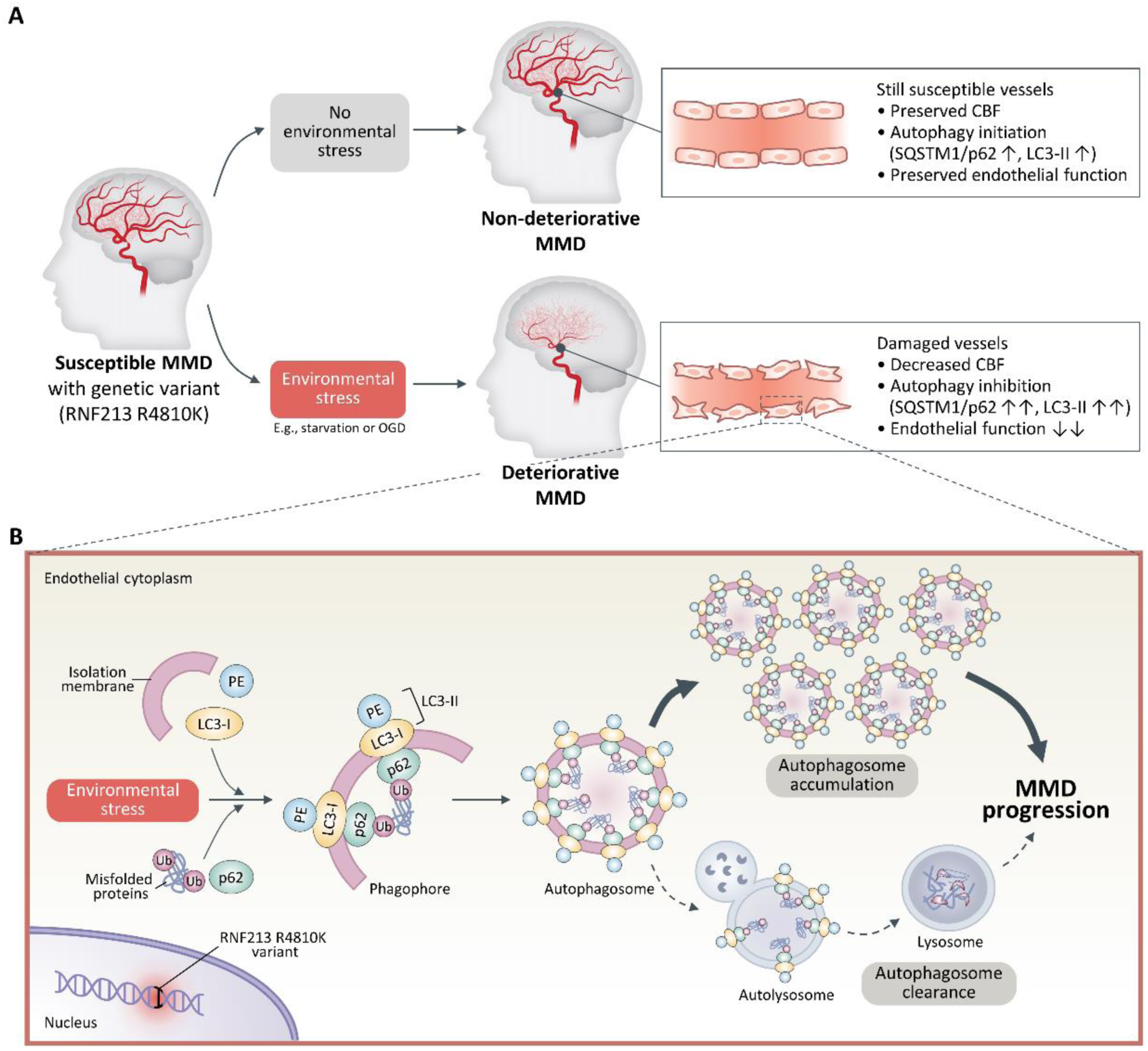
Endothelial dysfunction in MMD due to environmental stress: A potential mechanism of impaired autophagy in the RNF213 Variant. Possible mechanisms of MMD progression with the RNF213 R4810K variant under environmental stress. Upon susceptible MMD patients with the RNF213 R4810K variant, so-called non-deteriorative MMD, the disease phenotype tends to remain stable as long as there is an adequate blood supply to maintain cerebral blood flow (CBF). This stability is attributed to the effective clearance of cellular waste products, which are crucial for cellular homeostasis and mediated by autophagy. However, under certain environmental stressors, such as starvation or oxygen-glucose deprivation (OGD), the susceptible MMD patient experiences clinical deterioration, transitioning into what is named “deteriorative MMD.” Autophagic activity is suppressed under external stress. This inhibition of autophagy results in the accumulation of unnecessary proteins within endothelial cells. As these proteins accumulate, cellular homeostasis is disrupted, leading to endothelial dysfunction and the further progression of MMD.

### Damaged endothelial cells with RNF213 R4810K variant

Our findings revealed that the patient with MMD carrying the RNF213 R4810K variant had endothelial dysfunction as assessed by FMD. In addition, we observed that HUVEC^R4810K^ had reduced ability to form tubular structures when the cells were deprived of oxygen and glucose.

In clinical settings, patients diagnosed with MMD manifest various forms of endothelial dysfunction, such as thickening of the inner layer of blood vessels (intimal thickening) and disruption of the blood-brain barrier (BBB). ^33, 34^ Previous studies have also reported the presence of narrowed or steno-occlusive lesions in the cerebral proximal anterior circulation in MMD patients carrying the RNF213 variant. ^16, 35^ Endothelial function is commonly assessed using FMD, a non-invasive method through increased shear stress in response to reactive hyperemia. ^36, 37^ Numerous studies have reported that MMD patients are more prone to endothelial dysfunction, often tied to inflammation in the vessel walls. ^38^ This is in line with other findings where reduced FMD was observed in patients with acute ischemic stroke. ^39, 40^

Our study contributes to this body of knowledge by examining cells derived from induced pluripotent stem cells (iPSCs) of MMD patients with the RNF213 variant. These cells exhibited noticeable functional damage compared with similar cells from MMD patients with wild-type RNF213 wild type (WT). ^12^ In preclinical studies involving endothelial cell-specific overexpression of the RNF213 variant in mice, overexpression of the RNF213 variant led to critical problems in endothelial function and the inhibition of cerebral angiogenesis in the brain under hypoxic conditions. ^31^ Our dataset is consistent with that of previous studies indicating that impaired endothelial function plays a role in the constriction of blood vessels in the brains of patients with MMD. Additionally, our results provide a cellular model that imitates the decrease in blood flow and oxygen levels typically seen in the brains of patients with MMD.

### Induction of autophagy impairment by RNF213 R4810K variant

In this study, we observed that both patient-derived PBMCs and HUVECs carrying the RNF213 R4810K variant were more susceptible to autophagy impairment during OGD. There may be a link between this vulnerability and the fact that MMD-associated variants such as RNF213 R4810K cluster primarily within the ubiquitin-E3 ligase domain. ^20^ Previous studies have shown that this specific variant, the most prevalent single nucleotide polymorphism (SNP) in patients with MMD, is associated with defects in the ubiquitination process, a key cellular mechanism for protein degradation. ^20, 21, 41^

Our findings further confirm that this specific genetic variant leads to problems with autophagy, which is crucial for cellular self-maintenance and repair. Our data indicated that the RNF213 R4810K variant is an MMD susceptibility gene that causes impairment of autophagy. ^42, 43^ Recently, many studies have shown that autophagy in endothelial cells is essential for the recovery and regeneration of damaged tissue in the brain. ^44^ Notably, pathological processes are initiated when oxygen and nutrients are depleted. ^45^ Inflammatory signals or ROS generated under ischemic conditions are reduced by the activation of endothelial cell autophagy, thereby inhibiting endothelial damage. ^46^ However, activating autophagy in these cells helps minimize such damage, particularly by reducing harmful substances, such as ROS, and lowering inflammation caused by ischemic or hypoxic conditions. Therefore, our results suggest that enhancing or fine-tuning autophagy in endothelial cells may be an effective strategy for mitigating or even halting the progression of MMD.

### MMD treatment strategy using autophagy modulators

In this study, we demonstrated that CSZ could restore endothelial cell function via autophagy activation in HUVECs with the RNF213 R4810K variant under OGD conditions. Notably, our findings present a novel insight, as no previous studies have specifically investigated the role of autophagy modulation in the treatment of MMD.

The role of cilostazol in activating autophagy has been increasingly recognized in various medical fields. As a phosphodiesterase 3 (PDE3) inhibitor, CSZ was shown to significantly reduce mortality rates in patients with MMD compared with other treatment regimens involving clopidogrel, statins, or aspirin, according to a nationwide study. ^8^ Moreover, CSZ treatment has been associated with the prevention of the progressive narrowing of large intracranial arteries in cases of adult-onset MMD. ^45^

In preclinical settings, cilostazol effectively enhanced neuroprotection by activating autophagy in a rat model for Parkinson’s Disease induced by rotenone. ^47^ Similarly, CSZ demonstrates a protective role through autophagy activation in N2a cells, which are neuronal cells, in the context of Alzheimer’s Disease. ^48, 49^ In light of these observations, which align well with existing literature, we propose that our data may serve as fundamental evidence for a novel therapeutic strategy for MMD.

## Conclusions

Our research suggests that the process of autophagy plays a crucial role in the development of progressive endothelial dysfunction, leading to MMD in genetically susceptible patients who have the RNF213 R4810K variant. This indicates that autophagy is a critical regulator of endothelial cells, and dysfunction in this process could be a progressive cause of MMD. Our findings suggest that therapeutic approaches targeting autophagy modulation could be beneficial. Potential agents that restore impaired endothelial function and reactivate autophagy may offer a promising avenue for treating MMD.

### Novelty and Significance

#### What Is Known?

- Ring Finger Protein 213 (RNF213) is closely linked to Moyamoay disease (MMD) and has been identified as a key susceptibility gene.
- The RNF213 R4810K variant is induced in response to inflammatory, angiogenic, and hypoxic stresses, causing cellular dysfunction. Moreover, RNF213 R4810K variant exhibits a defect in ubiquitination.
- For patients diagnosed with diabetes mellitus (DM) and atherosclerosis, a notable impairment in autophagy is observed in their endothelial cells.

#### What Is New?

- An MMD patient aged 32, carrying the RNF213 R4810K variant, experienced rapid clinical deterioration coupled with marked shrinkage of intracranial vessels during a brief malnutrition episode.
- Autophagy inhibition is evident in patient-derived peripheral blood mononuclear cells (PBMCs) and human umbilical vein endothelial cells (HUVECs) expressing the RNF213 R4810K variant after exposure to oxygen-glucose deprivation (OGD). Despite being under OGD conditions, autophagy modulators (rapamycin and cilostazol) effectively restored the autophagic activity and function of HUVEC^R4810K^.

## Non-standard abbreviations and acronyms

MMD: Moyamoya disease
RNF213: Ring Finger Protein 213
OGD: Oxygen-glucose deprivation
PBMCs: Peripheral blood mononuclear cells
HUVECs: Human umbilical vein endothelial cells
FMD: Flow mediated dilation
TEM: Transmission electron microscopy
RA: Rapamycin
CSZ: Cilostazol

## Acknowledgements

Hee Sun Shin wrote the manuscript and was involved in the conception and design of this study, interpretation of data. Geun Hwa Park was involved in the conception and interpreted the data. Eun Sil Choi and Da Sol Kim interpreted the data. So Young Park was involved in acquisition of data. Dr Chang was involved in the critical revision for important intellectual content. Dr Hong was involved in the conception and design of this study, and interpretation of data, and critical revision for important intellectual content.

